# Striatal dopamine release reflects a domain-general prediction error

**DOI:** 10.1101/2023.08.19.553959

**Authors:** Kauê Machado Costa, Nishika Raheja, Jash Mirani, Courtney Sercander, Geoffrey Schoenbaum

## Abstract

Dopamine is classically thought to drive learning based on errors in the prediction of rewards and punishments ^1^. However, animals also learn to predict cues with no intrinsic value ^2^, and it is unclear if such latent learning also relies on dopaminergic prediction errors. Here, we tested this by recording dopamine release in the nucleus accumbens and dorsomedial striatum while rats executed a sensory preconditioning task that incorporates both types of learning ^3^. We found that dopamine release in both regions correlated with errors in predicting value-neutral cues during latent learning and with errors in predicting reward during reward-based conditioning. Moreover, dopamine in the nucleus accumbens reflected inferred value in the probe test, supported by orbitofrontal cortex activity. Our findings suggest that dopamine signals a domain-general, multi-factorial prediction error, capable of supporting model-based learning.

## Introduction

When navigating the world, animals learn both to predict rewards and punishments (value-based learning) and the relational structure of different elements of the environment, even if they are seemingly irrelevant (value-neutral or latent learning) ^2^. The first form of learning can be explained by a “model-free” framework, where learning is based on cached values, while the second form requires internal representations of the external world, or a “model-based” framework. The neuromodulator dopamine has been strongly implicated in value-based learning, as across several species and tasks dopamine activity reflects the difference between experienced and expected value, or reward prediction errors (RPEs) ^1,4–6^. This has led to the powerful and influential proposal that dopamine supports something akin to a model-free temporal difference reinforcement learning (TDRL) process in the brain ^1^.

However, recent work has questioned this interpretation. Dopamine signals have been found to correlate with, and influence learning about, variables that are not strictly value, including movement probability, interval timing, and sensory stimulus properties ^7–10^. Dopamine neuron activity also signals errors in predictions derived from value that must be inferred from task models ^11,12^. Some of this functional diversity has been related to differences in projection-specific subpopulations ^13,14^. For example, dopamine release in dorsal striatal regions, like the dorsomedial striatum (DMS), has been shown to correlate more with value-orthogonal variables than dopamine release in the nucleus accumbens core (NAcc) ^15–17^. However, this functional diversity is not sufficient to explain all of the data, since even within the NAcc, dopamine release can correlate with variables that are orthogonal to value, like novelty and sensory salience ^18,19^, and individual RPE-encoding dopamine neurons in the midbrain also signal errors in the prediction of task-irrelevant sensory properties of rewards, or sensory prediction-errors (SPEs) ^8,20^. Finally, optogenetic manipulations that causally affect RPE-based learning also impact learning about the sensory features of rewards ^21–23^.

To conciliate these findings with classical work, several new formal hypotheses on dopamine function have been proposed. These posit that dopamine neuron activity keeps tracks RPEs based on separate predictive “threads”, “bases”, or “channels”, resulting in RPE-signals that are dissociable according to their defining sensory or task-based properties ^24–26^. These models are essentially modifications of the current model-free framework to allow more complexity, diversity, or specificity in the RPE signal, however they remain tied to learning about motivationally significant elements.

Therefore, while these updated models can explain new findings on dopaminergic errors in learning about specific reward properties and the cues and actions associated with them, they do not predict any role for such error signaling in learning to predict events in the environment that are not related to reward or punishment.

Yet, such latent learning is common, and there is evidence that it can be influenced by brief activation and inhibition of midbrain dopamine neurons ^23^. While these results can be potentially explained by a permissive role for phasic or even tonic dopamine in value- neutral learning ^27,28^, it is also possible that they are evidence that dopamine operates as an error signaling mechanism that is *domain-general* or *multi-factorial*, which represents predictive errors across multiple dimensions, one of which is value^8,18,20,29–33^.

Here, we used optophysiological dopamine recordings and a classical learning theory task ^3^ to assess whether dopamine provides an error-like signal during value-neutral or latent learning. We recorded striatal dopamine signals while rats executed a 3-phase sensory preconditioning task (SPC), which incorporates value-neutral, explicit value-based, and inferred value-based prediction errors in its structure ^11,34^. We also took advantage of the known role of the lateral orbitofrontal cortex (lOFC) in supporting model-based inference in this task and inactivated this region during the critical probe session while recording dopamine signals ^34^. We found that dopamine release in both NAcc and DMS correlated with sensory prediction errors during the formation of the valueless cue-cue associations and with classical RPEs during conditioning. In the probe session, lOFC inactivation had a selective effect on NAcc dopamine release consistent with disruption of an inference process. These findings suggest that striatal dopamine signals reflect errors across different types of predictions, including when value is not at issue.

## Results

We transfected male rats with dLight1.2 and implanted them with optic fiber cannulas in the NAcc and DMS to allow simultaneous multi-site optophysiological recordings of dopamine release dynamics (Figure 1A and S1A) ^35–38^. To allow for better comparisons across rats and sessions, z-scored dopamine-dependent fluorescence was normalized to the maximum signal observed in each session (see Methods). We also transfected the same rats with either hM4d (inhibitory DREADD receptor; n=8; OFCi group) or mCherry (control; n=6; CTRL group) in the lOFC to allow us to chemogenetically inhibit activity in this region (Figure 1B) ^39,40^. After at least 4 weeks for recovery and viral expression, rats were food restricted, shaped to retrieve food-pellets from the food port, and then subjected to SPC training (Figure 1C). Dopamine release was recorded during all sessions, and JHU37160 dihydrochloride (JH60; i.p. 0.2 mg/kg), a high-potency DREADD agonist ^41^, was injected prior to the probe test to inactivate the principal neurons in the lOFC of the OFCi rats, as validated previously ^39,40^. Against this behavioral backdrop, we analyzed dopamine release dynamics in each recording site to address:

1. whether dopamine release in the probed locations reflected error-driven associative learning in the absence of overt value or reward,
2. if release in these same sites exhibited classic RPE signals during conditioning, and
3. how dopamine release was structured during the probe test, when the preconditioned cues were inferred to have value, a property known to depend on processing in lOFC.

**Figure 1.**
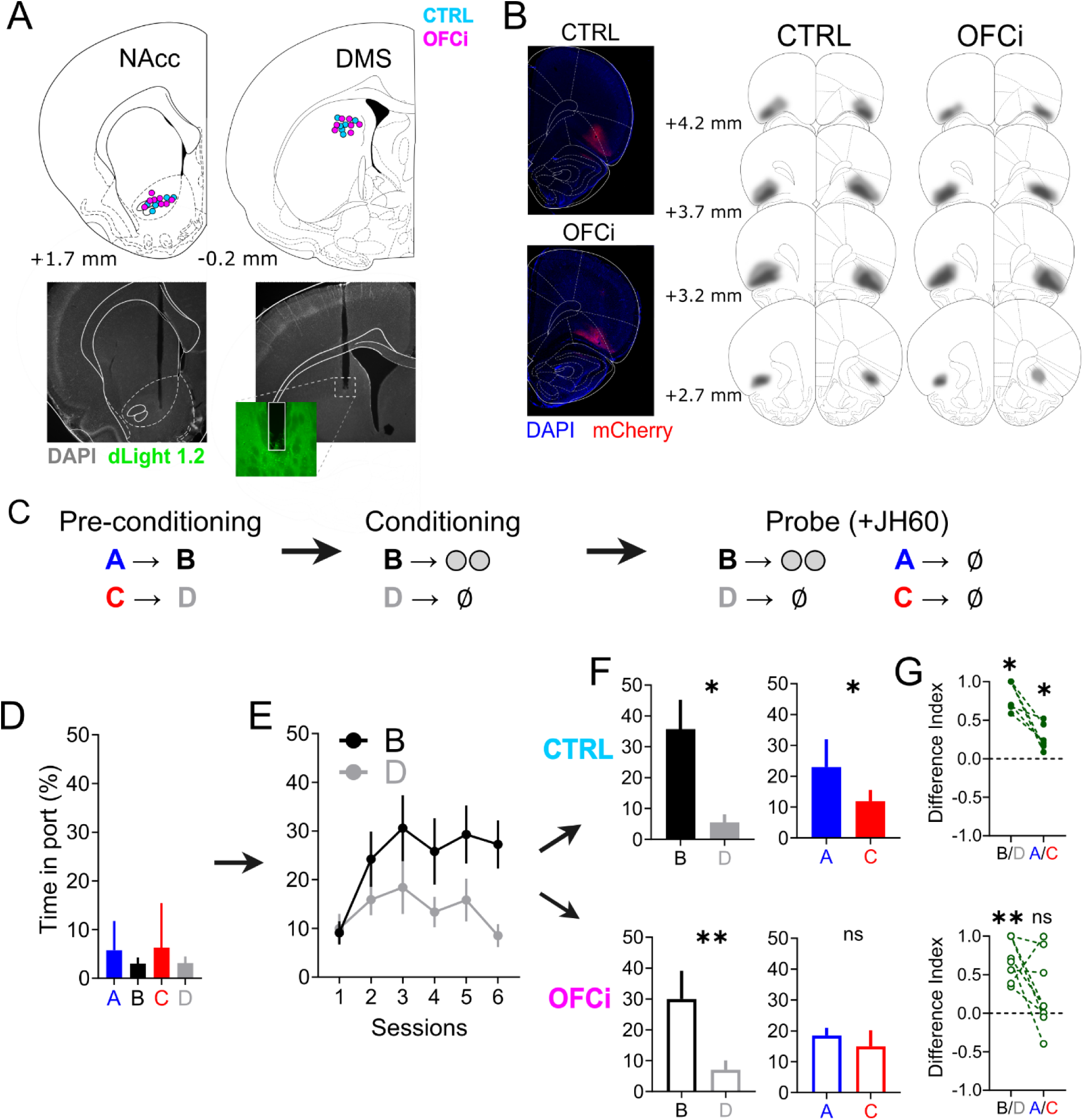
Histological verification and behavioral results. **A:** Location of fiber tips in the striatum for all recorded rats and striatal locations (top panels) and representative examples (bottom panels). Insert in the middle bottom panel contains a representative demonstration of the expression of dLight1.2 in relation to fiber tip location. **B:** Spread of DREADD construct expression (mCherry reporter) in lOFC. Left panels contains representative histological microphotographs of a CTRL and an OFCi rat. Right panels contain schematic reconstructions of the viral spread across the lOFC in both groups. Lighter shading represents area of maximal spread, while darker shading reflects the area of minimal spread. **C:** Schematic of the SPC behavioral task. **D:** Behavioral responding, as measured by total time spent in the food port during cue presentation, for all rats in the preconditioning phase. Due an absence of group effects (CTRL vs OFCi) in the preconditioning and conditioning phases, these data were pooled together in this figure for clarity (see Figure S2 for results split according to groups). **E:** Behavioral responding in the conditioning phase. Note the progressively increased responding to the reinforced cue B in relation to the nonreinforced cue D. **F:** Behavioral responding during the probe test, where all rats were injected with the DREADD agonist JH60. **G:** Difference index (difference divided by the sum) between responding to cues B and D and cues A and C, for the CTRL and OFCi groups. A positive difference index indicates a higher responding to B or A, while a negative one indicates higher responding to C or D, in each of the comparisons. Difference index values were tested with one-sample Wilcoxon tests against a hypothetical 0 (equal responding to both cues). This is essentially another way of representing the data from the graphs in F and it confirmed a significant higher responding for B versus D in both groups (*P*=0.031* for CTRL and *P*=0.008** for OFCi) and a similar higher responding for A only in the CTRL group (*P*=0.031*) and not in the OFCi group (*P*=0.148). Note from F and G that rats in the CTRL group, where OFC activity was not affected, respond more to cue A, which in preconditioning predicted the subsequently reinforced cue B, than to cue C, which was paired with the subsequently unreinforced cue D. In the OFCi group, where OFC activity was chemogenetically inhibited, there was no difference in responding to cues A and C. Also notably, rats in both groups responded significantly more to B than to D, irrespective of OFC inactivation. Data are represented as mean ± SEM. \**P*<0.05; \*\**P*<0.01.

### Behavioral results and the role of lOFC in inference-based behavior

First, we consider the behavioral results. During the preconditioning phase, food port responding during the presentation of each cue was low and rats did not distinguish between them (Figure 1D; 2-way ANOVA for cue and group effects: *P*>0.113 and *F*<2.131 for all comparisons). During conditioning, responding progressively and selectively increased to B, the cue paired with food delivery (Figure 1E; 3-way ANOVA for effects of session progression, cues, and group; session effect: *P*=0.014*, F_(5,60)_=3.152; cue effect: *P*=0.002**, F_(1,12)_=14.88; interaction of cue and session effects: session effect: *P*=0.008**, F_(5,60)_=3.497; all other effects: *P*>0.742, *F*<0.327).

Subsequently, in the probe test, all rats responded more to B, the food-paired cue, than to the non-reinforced cue, D, indicating that they had correctly learned to attribute higher value to the food-paired cue (Figure 1F; Wilcoxon test, *P*=0.031* for CTRL and *P*=0.008** for OFCi). Furthermore, rats in the CTRL group responded more during A than C (*P*=0.031*), indicating that they had learnt the A-B association in the pre- conditioning phase and were able to use it in the probe test to infer that A (and not C) might also lead to reward (as it predicted B). Rats in the OFCi group, on the other hand, had similar responding to cues A and C (*P*=0.313), despite showing normal B versus D discrimination, indicating a selective disruption of inference-guided behavior, replicating previous results obtained with other methods of lOFC inactivation ^34,42^.

### Dopamine release in the NAcc and DMS exhibit error-like correlates during pre- conditioning

We next asked how dopamine release responded during the pre-conditioning phase, when the rats were exposed to arbitrary pairings of neutral cues, in the absence of any value, overt reward, or even response requirement. Focusing on release around cue presentation, and starting with responses in the NAcc, we observed clear dopamine peaks at the onset of each cue (A,B,C, and D). Initially, these peaks were similar (Figure 2A and 2B) and each diminished with repeated exposure (2-way ANOVA for effect of trial progression and cue predictability; trial effect: *P*<0.0001***, *F*_(2,54)_ = 22.76), a pattern consistent with signaling of surprise or novelty ^43–45^ and which has been associated with learning that underlies latent inhibition ^19,46^.

**Figure 2.**
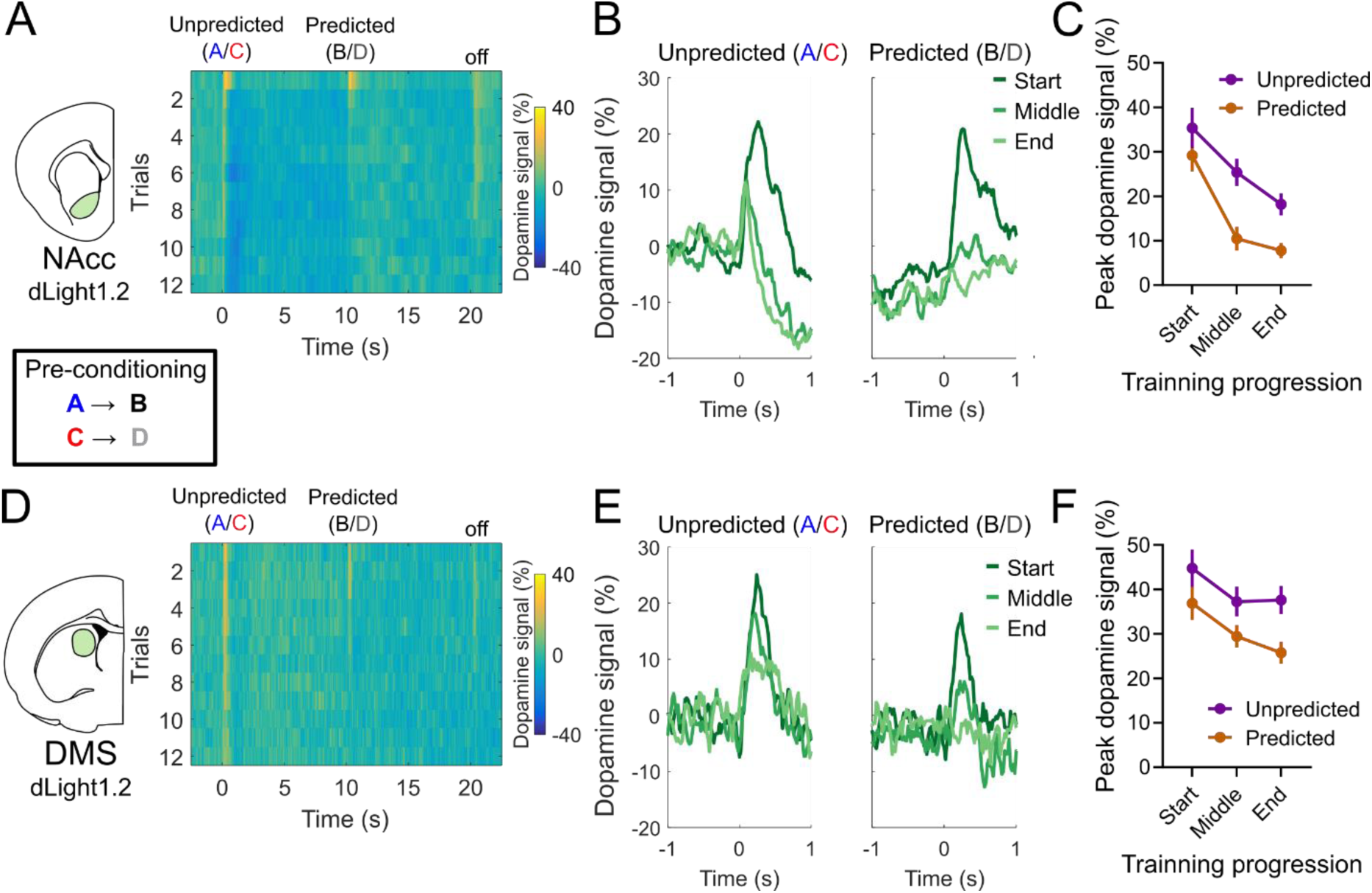
Dopamine signals in NAcc and DMS reflect general prediction errors during sensory preconditioning. **A:** Heatmap representation of average NAcc dopamine responses during all 12 preconditioning trials, recorded with dLight1.2 (N=14). Note the larger dopamine responses during the first trials to both the unpredicted and predicted cues, and the progressive diminishment of the responses with subsequent trials, which is more pronounced for the responses to predicted stimuli. **B:**.Average traces of NAcc dopamine responses to predicted and unpredicted cues during the start (first two trials), middle (trials 6 and 7), and end (trials 11 and 12) of the preconditioning phase. **C:** Peak NAcc dopamine responses measured over the course of the preconditioning training. Note the similar responses to both types of cues in the beginning of training, and the significant difference between responses to predicted and unpredicted cues. **D-E:** similar to A-C, but for DMS dopamine signals, simultaneously recorded with NAcc signals. We highlight that in the DMS, the progressive reduction of dopamine responses to both types of cues was not as strong as in the NAcc. Data are represented as mean ± SEM. \**P*<0.05; \*\**P*<0.01.

However, concurrently with this expected effect, we also found that peak responses to the predicted events, B and D, diminished faster and to a greater extent than those to the unpredicted stimuli, A and C (Figure 2B and C; cue predictability effect: *P*=0.0004**, *F*_(1,27)_ = 16.48). This additional effect indicates that the predictive structure influences the dopamine responses to each cue, essentially providing an error-like correlate during value-less sensory-sensory learning – a SPE riding on top of novelty-related changes (Figure 2).

We observed a qualitatively similar response pattern in the DMS, where dopamine peak responses were larger at the first presentation of each cue, reduced over repeated exposures, and were lower for predicted events than for unpredicted events (Figure 2D and 2E; trial effect: *P*=0.023*, *F*_(2,54)_ = 4.039; cue predictability effect: *P*=0.004**, *F*_(1,27)_ = 9.577). However, the changes in peak responses over time were less pronounced and seemed to occur at a slower progression rate than in the NAcc, indicating that perhaps the changes observed in DMS responses were following those in the NAcc, consistent with previous evidence suggesting an ascending ventromedial to dorsolateral hierarchy of learning-induced plasticity in midbrain-striatal circuits ^47,48^.

To confirm the importance of the predictive structure in determining the changes in dopamine release that we observed, we trained another group of rats (n=5), this time transfected with GRAB-DA2m and implanted with an optic fiber cannula only in NAcc and without viral injection of DREADDs into lOFC. These rats underwent the same pre- conditioning training as above (Figure 3A-C), except that we added a session at the end in which we switched the order in which the cues were presented (Figure 3D). In other words, in this added session, the rats initially received presentations of A→B and C→D, as they had the prior two sessions, followed by presentations of B→A and D→C. If the changes in dopamine release were due to the predictive structure of the cues, then the release should be immediately restored to D and B when they appear unpredicted by another cue, and it should slowly diminish to A and C, as the new predictive structure is learnt. Consistent with this idea, we first found that VMS dopamine signals in this new group showed similar SPE signature as in the previous dLight1.2-transfected rats during the first 2 sessions - responses to predicted cues B and D were lower than responses to the unpredicted cues A and C - confirming that this finding is robust across experimental groups and dopamine sensors (Figure 3A-C; trial effect: *P*=0.0002***, *F*_(2,18)_ = 14.86; cue predictability effect: *P*=0.008**, *F*_(1,9)_ = 11.43). Then, when the order of cue presentation was reversed in the second half of the added preconditioning session, the responses to B and D, now unpredicted, were restored, and responses to cues A and C, now predicted, were slightly diminished (Figure 3D-F; 2-way ANOVA for effect of cue predictability – unpredicted versus predicted - and cue identity – B/D versus A/C; predictability effect: *P*=0.0228*, *F*_(1,9)_ = 7.512; cue effect: *P*=0.3102, *F*_(1,9)_ = 1.156;). This pattern strongly suggests that VMS dopamine release reflects the predictive structure of cue presentation in exactly the same way that the classic RPE response reflects the order of presentation of a cue and an event with overt value – an SPE for lack of a better term.

**Figure 3.**
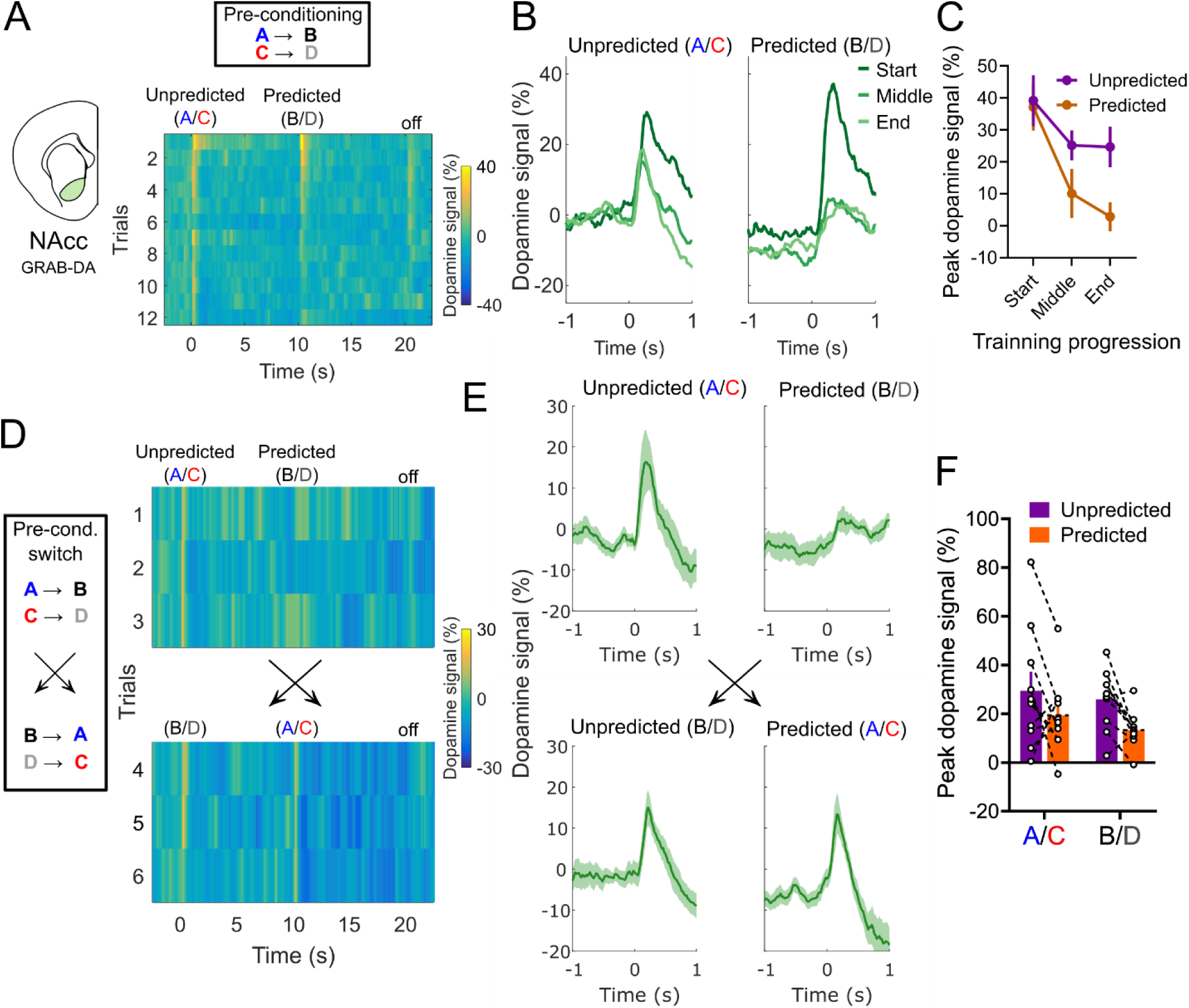
Encoding of sensory-sensory prediction errors by dopamine signals in NAcc depends on the predictive relationship between cues. **A:** Heatmap representation of the average NAcc dopamine signals during all trials of preconditioning training in a separate cohort of rats (N=5), recorded with GRAB-DA2m. Note that the response pattern is similar to those recorded with dLight1.2. **B:** Average traces of dopamine responses during the start, middle, and end of preconditioning, defined the same way as in Figure 2. **C:** Peak dopamine responses during preconditioning training in this cohort of rats. Note the similar pattern to the cohort in Figure 2. **D:** Heatmap representation of dopamine signals in the preconditioning switch session. In the first three trials the structure of the task was the same as in the previous training sessions, with A preceding B and C preceding D. In the last three trials, that order was switched, with B not coming before A and D before C, with no external signaling to the rats. Note that responses to B and D, which were almost entirely suppressed in trial 3, were immediately restored in the trial 4, when these cues presented unexpectedly, and responses to cues A and C, which were unexpected, start to become slightly more suppressed once they become predicted by cues B and D. **E:** Average dopamine responses to each type of cue before and after the switch. **F:** Peak dopamine responses to cues A/C and B/D when they were both predicted (first three trials for B/D and last three trials for A/C) and unpredicted (first three trials for A/C and last three trials for B/D). For both sets of cues, regardless of their identity or exposure history, NAcc dopamine responses were larger when they were unpredicted than when they were predicted. Data are represented as mean ± SEM.

### Dopamine signals in NAcc reflect classical reward prediction errors during conditioning

Consistent with numerous prior studies, we found that peaks in dopamine release in the NAcc strongly signaled RPEs during the conditioning phase of the task (Figure 2)^15,17^. Peaks in release were initially high in response to pellet delivery and then decreased over time, while peaks in response to onset of the pellet-paired cue, B, increased (Figure 4A). This pattern could be resolved on a trial-by-trial (Figure 4A) or session-by-session basis (Figure 4B), with the difference between peak responses to the pellet delivery (US) and the cue (CS) becoming progressively lower with training (Figure 4C; 2-way ANOVA for effects of session progression and cue – B versus D; session effect: *P*<0.0001***, *F*_(5,65)_ = 14.48; cue effect: *P*<0.0001***, *F*_(1,13)_ = 153.2; interaction effect: *P*<0.0001***, *F*_(5,65)_ = 9.809)^11^. Importantly, responses to the non-reinforced cue, D, did not change significantly over time. These results confirm that dopamine signals in NAcc - recorded in the current rats, fiber locations, and using this task and methods - exhibited classic RPE correlates during conditioning.

**Figure 4.**
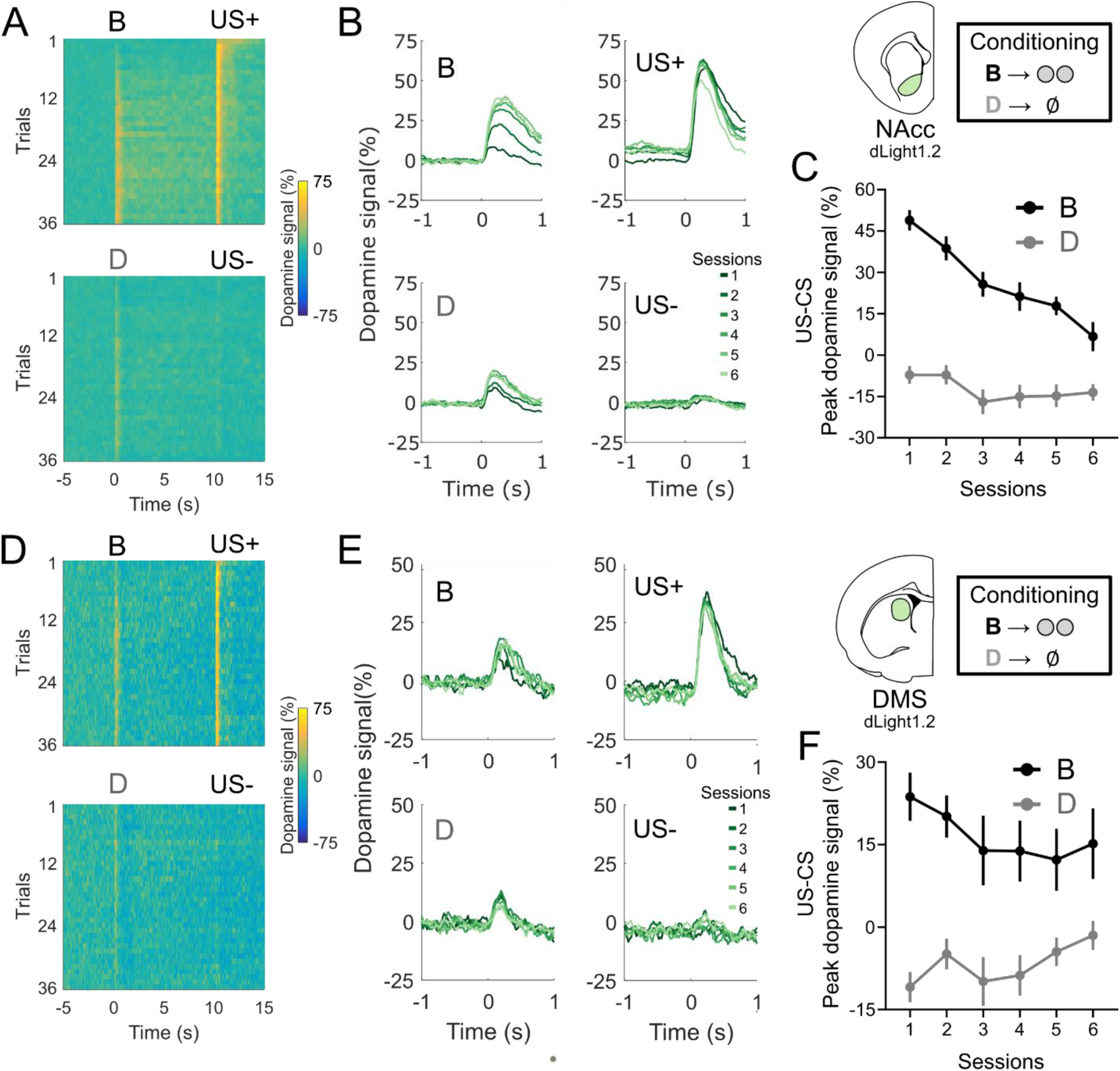
Dopamine release in VMS and DMS reflects classic, value-based, “reward” prediction errors in the conditioning. **A:** Heatmap representation of average NAcc dopamine signals in all trials of the conditioning phase of the cohort recorded with dLight1.2 (N=14). Note the progressive reduction of responses to US presentation and concurrent increase is responses to cue B, the reinforced CS+, with less changes in the responses to cue D, the unreinforced CS-. **B:** Average traces of dopamine signals for each session (6 trials per condition per session) of conditioning. **C:** Difference between the peak NAcc dopamine responses to US and CS presentation, for both cues B and D. A progressive decrease in this difference is key indicator of an RPE response ^11^. **D-F:** Same as A-C, but for DMS dopamine signals. Note that DMS dopamine also shows signs of RPE encoding, but much less prominently as NAcc. Data are represented as mean ± SEM. \**P*<0.05; \*\**P*<0.01.

Furthermore, we found that dopamine peaks in the DMS also seemed to reflect RPEs, albeit again with a lower update rate as compared to the NAcc, and with less of a difference in relation to the peaks to the CS- (Figure 4D-F; session effect: *P*=0.492 *F*_(5,65)_ = 0.892; cue effect: *P*<0.0001***, *F*_(1,13)_ = 36.26; interaction effect: *P*=0.0221*, *F*_(5,65)_ = 2.842). This parallels our findings in the preconditioning phase, where changes to signals in DMS seem to lag changes to signals in NAcc.

### Dopamine release in the NAcc and DLS responses to cues with inferred value are dependent on lOFC activity

Finally, we arrive at the probe phase of the main SPC experiment. Up until now we have collapsed the two groups because there was no effect of group on any of the previous comparisons (Figures S2 and S3; 3-way ANOVA for effects of session/trial progression, cues, and group; all group main or interaction effects: *P*>0.1, *F*<0.5). In the NAcc, as expected, responses to the reinforced cue B were significantly higher than to the unreinforced cue D in both the CTRL and OFCi groups (Figure 5A-B; paired t-tests; CTRL: P= 0.002**; OFCi: *P*=0.007**). In the DMS, there was no significant difference between responses to B and D cues in either group (Figure 5A-B; paired t-tests; CTRL: P= 0.087; OFCi: *P*=0.466), which is likely reflective of the diminished RPE encoding observed in DMS and the lower number of trials in the probe session.

**Figure 5.**
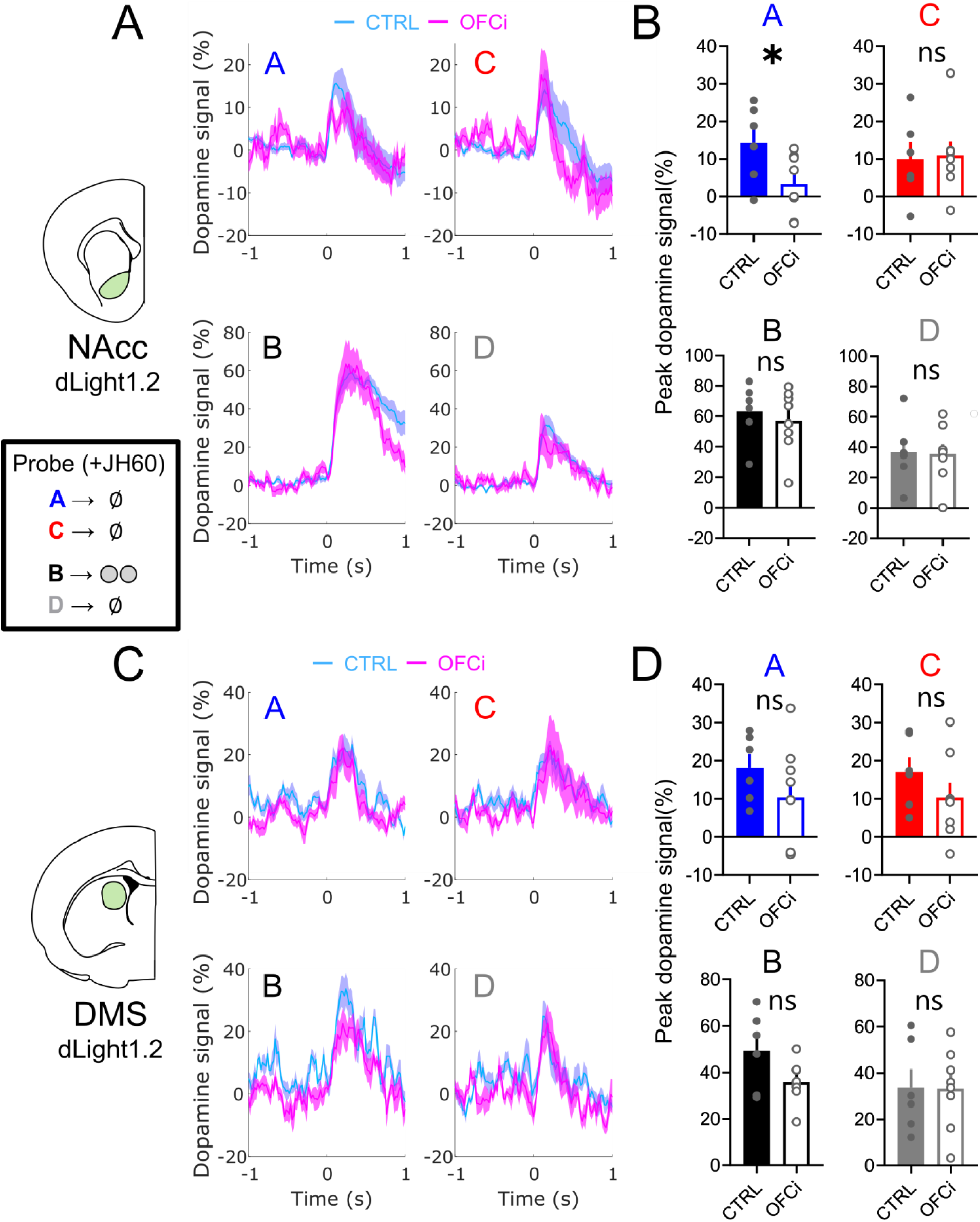
Dopamine signals in NAcc and DLS reveal a contribution of lOFC on learning-dependent dopamine signals during probe. **A:** Average traces of NAcc dopamine responses to all cues in the probe test, for both CTRL (N=6) and OFCi (N=8) rats. All recordings and behavior in this session were performed after the i.p. injection of 0.2 mg/kg of JH60, which inhibited OFC activity in the OFCi group. Note that the only difference between responses in the two groups is a blunted dopamine peak to A in the OFCi group. **B:** Peak NAcc dopamine responses to all cues in the probe session, confirming the significant difference only in responses to A between CTRL and OFCi rats. **C-D:** same as in A-B, but for DMS dopamine responses. In the DMS, there was no significant difference between responses to any of the cues between the CTRL and OFCi rats. Data are represented as mean ± SEM. \**P*<0.05.

Surprisingly, in the CTRL group dopamine responses to cues A and C were similar during the probe phase (Figure 5). This was true in both areas (paired t-test, A vs C in CTRL group; NAcc: *P*=0.326; DMS: *P*=0.801). That measured dopamine release did not differ for the two cues was unexpected, particularly in light of the differential behavior of these rats to the two cues in the probe test (Figure 1) as well as prior unit recording data showing that dopamine neuron spiking is higher to cue A than to cue C ^11^. These results are consistent with prior demonstrations that striatal dopamine release and dopamine neuron firing are not always correlated, and that certain behavioral variables can be observed in one measure but not the other ^49^.

Nevertheless, even if the release appears similar, the hypothesized circuit function predicts that release should reflect different information for the two different cues; specifically, while release to cue C might reflect non-specific generalization and perhaps elevated salience due to the long period since it was last seen, release to cue A should reflect, at least in part, the specific predictive relationship between A and B and between B and reward. We know from prior work in this setting that this is the case for the behavior to A, as it is sensitive to devaluation of the predicted reward ^50^. Critically, behavior based on this information is selectively sensitive to inactivation of lOFC ^34^. This predicts that inactivation of lOFC should selectively affect dopamine release to A .

Consistent with this prediction, we found that OFC inactivation did reduce the relative peak responses to cue A in NAcc (Figure 5A-B), with responses to all other cues being relatively similar (paired t-tests; A vs C in OFCi group; NAcc: *P*=0.044*) (unpaired t- tests; A response in CTRL vs OFCi; NAcc: *P*=0.043*; all other comparisons of CTRL vs OFCi showed *P*>0.05 for both NAcc and DMS).

## Discussion

Here we recorded dopamine in a three-phase SPC task and found that release in both NAcc and DMS reflected errors in predicting neutral cues during the initial preconditioning phase of training. These signals were dissociable from sensory salience and detected at sites that also showed classic RPE signals during subsequent conditioning. Thus, the results replicate previous findings linking striatal dopamine release to salience and signaling of value-based prediction errors ^1,15,18,19,44^, while also showing a novel, value-independent error-like response during learning about neutral cue pairs. The significance of the latter finding is bolstered by previous work using optogenetics showing that dopamine activity is both necessary and sufficient for driving sensory-sensory learning in a similar preconditioning task ^23^. The causal manipulations in this previous work could not differentiate between an error-driven signal and a more general permissive role of dopamine for value-neutral learning, thus the current data are critical in ruling out these possibilities in favor of a prediction error-like mechanism.

Importantly, showing error signals in this setting goes beyond prior work by us and others showing error signals during learning of specific features of otherwise valuable events (i.e. rewards) ^8,12,20,21,51^, which can be explained with adjustments to TDRL algorithms, such as the addition of dissociable “threads”, “bases”, or “channels” for keeping track of different components of rewarding events ^24–26^. These models cannot easily explain why neutral cues evoke error signals. Instead, the response patterns observed here to neutral cues are much better explained by a model-based process, or at least a hybrid model incorporating something like successor representations ^29,30,32,40^.

These results, coupled with much prior work ^1,8,12,17,20,21,33,51^, suggest that dopamine activity is shaped by multiple factors or dimensions that characterize events, of which value is but one component ^29,30,32,40^. This broader interpretation holds better explanatory power than several alternatives. For example, one potential challenge to our conclusion that dopamine release reflects more than value is the expansion of the concept of “value” itself. For example, novel sensory cues may be assigned intrinsic value, given that they could potentially convey information, about the external world ^52–54^, or perhaps previous experiences, in the home cage or during shaping, led the rats to acquire the general belief that sounds are broadly predictive of rewards prior to preconditioning ^55^. These explanations seem unlikely to us, since preconditioned cues trained in this manner do not support conditioned reinforcement ^56^. Further, if these cues were intrinsically valuable, like a primary reward, then the dopamine responses to the unpredicted cues A and C should have increased over time during preconditioning, just like responses to B during conditioning, but they did not. Instead, the observed pattern, in which dopamine responses to predicted cues were inhibited faster, seems more compatible with the formation of a predictive association between two cues without intrinsic value.

A related set of ideas is that dopamine responses track stimulus salience ^18,31,44^. This component of the signal differs from a value prediction error because it factors in attributes of the cues themselves and the relational structure of task elements, such as novelty, stimulus intensity, and unexpectedness. Its presence should increase associability or learning about a cue, both at that time and subsequently. Our findings here and in prior causal work are generally compatible with this interpretation, as predicted cues would indeed, by definition, be less surprising than unpredicted cues. However, contrary to predictions for salience, there was no impact of activating dopamine neurons on *subsequent* learning in our prior work ^23^.

Indeed, novelty, sensory salience, and signaled sensory and value prediction errors, could all be conceptualized as prediction errors that operate on different variables and across different hierarchical structures of time scale and information content ^8,32^. Similar frameworks have been proposed to shape perception and general cognition, and the idea that the brain itself is fundamentally dedicated to processing, and minimizing, prediction errors is currently an influential and powerful hypothesis ^57^.

The involvement of dopamine in model-based learning is also supported by the effects of lOFC inactivation in the probe test, where it selectively disrupted behavioral responding and dopamine release in NAcc to cue A, the cue endowed with inferred value by virtue of the prior training. This result is consistent with the impact of muscimol inactivation of lOFC on responding in the probe test in our prior work ^34^, and with the widespread observation that the lOFC is strongly associated with inferential or model-based behavior across a variety of tasks, modalities, and species ^58^.

Dopamine recordings in the probe test did show some unexpected features. In the control group, dopamine release was similar both to cue A, the preconditioned cue predictive of reward, and cue C, the control cue. If the only mechanism at work were one based on inference and if the only information represented by the error signal were value, then the signal should preferentially have shown up to cue A. In this regard, the results contrast with the results of a previous single unit recording study ^11^, in which the firing of putative dopamine neurons in the ventral midbrain differed between cues A and C in the probe test of a similar task.

However there may be a number of non-mutually exclusive explanations for this finding. For starters, the optophysiological methods used here may be not sensitive enough to pick up what may be a very subtle difference in the dopamine responses to cues A and C in the control group, or to put it another way, other components of the dopamine response (i.e., local features of the spatiotemporal patterns of release, differential release from individual axon terminals, etc.) may better reflect inferred prediction errors than bulk responses to cue onset. It is also possible that in the unit recording study ^11^ the sampled neuronal population may have contained a disproportionate number of neurons that projected to other striatal regions not sampled in our current work, where different variables may be represented. However, the methods and analyses applied here, which fit the standard of the field, were successful in detecting prediction errors in preconditioning and conditioning, including classically defined RPEs, as well as some differences imposed by lOFC inactivation. Therefore, this result seems unlikely to be due to these artifactual causes.

More likely, release may follow a different time course than spiking activity, with different sensitivities to aspects of our design. Such speculation is consistent with recent work showing that dopamine neuron firing and release in target regions can be dissociated, likely due to the sculpting effects of local striatal mechanisms ^49,59–61^. Similar release in controls to the two preconditioned cues might reflect a greater sensitivity of these release dynamics to commonalities across the cues that are unique to our task. For instance, in preconditioning, all the cues are initially presented without any reward, and this similarity may be the foremost determinant of bulk dopamine release signals in the probe, even if other processes are in play. This might be quite different for other tasks designed to study inferential behavior, like reinforcer devaluation, where no common unrewarded latent state can be inferred during the initial learning phase ^12^. Within this framework, one might predict that it would require a selective manipulation of areas that process inferential information – such as the lOFC - to reveal a potential difference in dopamine responses that depend on such processes, which is indeed what we found.

In conclusion, our findings support the proposal that dopamine signaling in the striatum mimics a *general*, *multi-factorial* prediction error term that comprises different variables and whose specific information content varies across different regions. These prediction errors in the NAcc and DMS reflect the relational structure of value-neutral sensory-sensory associations, as well as cached value-based associations and novelty. We also present evidence that accumbal dopamine responses to cues with inferred value are differentially sensitive to lOFC disruption, suggesting that dopamine signals that appear similar may be dependent on different underlying mechanisms according to their task relevance.

## Acknowledgements

We thank Marios Panayi, Angela Langdon, Matthew Gardner, and Evan Hart and for critical discussions. This work was supported by the Intramural Research Program at the National Institute on Drug Abuse. KMC received a post-doctoral fellowship from the German Research Foundation (Deutsche Forschungsgemeinschaft, DFG; MA 8509/1- 1).

## Author contributions

KMC and GS designed experiments, interpreted results, and wrote the manuscript. KMC performed the experiments with technical assistance from NR, JM, and CS. KMC analyzed the data and wrote the first draft of the manuscript. GS supervised the research.

## Materials and correspondence

All data and code displayed in this manuscript will be made available in a publicly accessible repository. Additional information on materials and protocols are available upon request to Geoffrey Schoenbaum.

## Methods

### Experimental Model and Subject Details

Experiments were performed on a total of 31 male Long-Evans rats (>3 months of age at the start of the experiment, Charles River Laboratories) housed on a 12 hr light/dark cycle at 25 °C. Rats were food restricted to ∼85% of their original weight for the duration of the experiments and were tested at the NIDA-IRP in accordance with NIH guidelines determined by the Animal Care and Use Committee, which approved all procedures. All rats had *ad libitum* access to water during the experiment and were fed 16-20 g of food per day, including rat chow and pellets consumed during the behavioral task. Behavior was performed during the light phase of the light/dark schedule. Prior to surgery and each experiment, rats were handled by the experimenter, as previously described ^39,40^.

### Surgical procedures

For the main preconditioning experiment, rats (N=24) were anesthetized with 1-2% isoflurane and prepared for aseptic surgery. They received unilateral infusions of AAV5- hSyn-dLight1.2 into the NAcc (AP +1.7 mm, ML + or -1.7 mm, and DV -6.3 and -6.2 mm from the brain surface), DMS (AP -0.4 mm, ML + or -2.6 mm, and DV -3.7 and -3.6 mm from the brain surface) and dorsolateral striatum (DLS, AP +0.2 mm, ML + or -3.8 mm, and DV -3.7 and -3.6 mm from the brain surface). Recordings from DLS were poorly correlated with our task events, and seemingly correlated more with individual movements; as that was not the focus of our study, these signals were not further included in our analysis. A separate group of rats received unilateral infusions of AAV9- hSyn-GRABDA2m (N=5) or AAV9-hSyn- GRAB-DA-mut (N=2) only in the NAcc, and no additional viral infusions. A total 0.7 μL of each dopamine sensor-transfecting virus was delivered in each site at 0.1 μL/min via an infusion pump. Optic fiber cannulas (200 μm diameter; Neurophotometrics, CA) were implanted in each site in the location of the second (most dorsal) viral infusion.

Rats for the main preconditioning experiment, which were transfected with dLight1.2 and implanted in different areas of the striatum, also received in the same surgery infusions of either AAV8-CaMKIIa-hM4d-mCherry (an inhibitory, Gi-coupled, designer receptor exclusively activated by designer drugs (DREADD); OFCi group, N=12) or AAV8-hSyn-mCherry (CTRL group, N=12), bilaterally into the lOFC (AP −3.0 mm, ML ± 3.2 mm, and DV -4.4 and -4.5 mm from the brain surface) ^39^. A total 0.5 μL of each DREADD virus was delivered in each site at 0.1 μL/min via an infusion pump. All viruses were obtained from Adgene. Exposed fiber ferrules and a protective black 3D-printed headcap were secured to the skull with dental cement. After surgery, rats were given Cephalexin (15 mg/kg po qd) for two weeks to prevent any infection. Two rats in the OFCi group and one rat in the CTRL group lost their headcaps after surgery and were removed from the study.

### Fiber photometry

Dopamine-dependent fluorescence signals were recorded using custom ordered multi-pronged fiber optic patch cables (200 μm diameter, 0.37 NA, Doric Lenses, Canada) that were attached to the optic fiber ferrules on the skull of the rats with brass sleeves (Thorlabs, NJ). Up to three fibers were connected at a time in each rat for recordings, and they were shielded and secured with a custom 3D-printed headcap-swivel shielding system that allowed for the relatively free movement of the rats without the use of optic commutators and prevented the spillover of light during recordings.

Recordings were conducted using an FP3002 system (Neurophotometrics, CA), by providing both 470 (active signal) and 415 nm (isosbestic reference) excitation light through the patch cord in interleaved LED pulses at 100 Hz (50 Hz acquisition rate for each channel). The light was reflected through a dichroic mirror and onto a 20× Olympus objective. Excitation power was measured at ∼150-200 µW at the tip of the patch cord. Emitted fluorescent light was captured thorough the same cords, separated with an image splitting filter, and captured via a high quantum efficiency CMOS camera. Signals were acquired and synchronized with behavioral events using Bonsai ^37^. To avoid photobleaching, recordings were triggered so that the system was only active during behavioral trials (see next section).

Signals were processed using custom scripts in MATLAB (MathWorks, MA). Raw fluorescence signals from the 470 (active) and 415 (reference) channels were first filtered with a fourth-order median filter. Then the reference channel data was fitted to the active signal using a second order polynomial fit, and the fitted data was subtracted from the active channel, removing dopamine-independent variations in fluorescence.

The resulting signal was z-scored for each trial, using the 10 seconds before each trial onset as a baseline. The resulting z-scores were normalized for each rat and session, were the maximum observed peak was set to 100% and the mean was set at 0%.

For quality control, a group of rats (n=2) were infused with GRAB-DA-mut, a mutated version of the GRAB-DA sensor that does not show dopamine-dependent fluorescence and tested in similar tasks as the other experimental groups. All recordings in this group showed no fluorescence peaks around any behavior event, confirming that the signals we observed in the other groups were indeed due to dopamine release (Figure S1 and S4).

### Behavioral apparatus and general procedures

Rats were trained and tested at least eight weeks after the surgeries. All experiments were conducted in standard behavioral boxes (12” x 10” x 12,” Coulbourn Instruments, PA), which were individually housed in light- and sound-attenuating boxes (Jim Garmon, JHU Psychology Machine Shop). Each box was equipped with a food cup (recessed magazine, a pellet dispenser, two wall speakers (one for white noise, and the other for tones and sirens), a clicker, a house light, and a panel light. Head entries into the food cup was measured based on breaks of an infra-red beam. For conditioning, 45 mg sucrose pellets (5TUT, TestDiet, MO) were used as reinforcers. Intertrial intervals (ITIs) varied around ∼6 min on average and all cues lasted 10 seconds. Each rat experienced one session per day, and the order of presentation of each cue was counterbalanced across rats. Behavioral responses were quantified as the percentage of time that each rat spent in the food cup during the last 5 seconds of each CS.

A computer equipped with GS3 software (Coulbourn Instruments, PA) controlled the equipment. One Arduino microcontroller in each box recorded all events from the Coulbourn equipment using a strobe based logic and relayed it as a serial input to the Bonsai photometry interface. One master Arduino relayed trial initiation and end from the Coulbourn interface to the FP3002 photometry hardware, triggering photometry recordings only during trials and turning them off during the ITIs.

### Sensory preconditioning task

Rats were first lightly shaped to retrieve sucrose pellets from the food cup. They were then submitted to two sensory preconditioning sessions; these used a total of four different auditory stimuli, drawn from stock equipment available from Coulbourn, which included tone, siren, clicker, and white noise. Assignment of these stimuli to task cues was counterbalanced across rats. In each pre-conditioning session, rats received six blocked presentations of A-B pairs (A presented unexpectedly, followed by the immediate presentation of B) and C-D pairs (similar arrangement as the A-B pairs), with no direct reinforcement being present.

After pre-conditioning, rats in the main experiment were conditioned for six sessions, where they received randomized presentations of B immediately followed by delivery of two sucrose pellets, and D without any reinforcement.

In the final probe session, rats experience three reminder trials each where B was presented followed by reward, and D was presented with no outcome, immediately followed by blocked presentations, six each, of A and C, with no outcome. Prior to this probe, each rat received an i.p. injection of JH60 (0.2 mg/kg, dissolved in 0.9% NaCl) and was left in their home cage for at least 20 minutes before the start of the session, to allow for the DREADD agonist to effectively inhibit transfected lOFC neurons in the OFCi group ^39,41^. Some rats in both groups did not show proper discriminative conditioning in both the conditioning and probe sessions. To control for this, only rats that responded twice as much to cue B than to cue D in the probe reminder trials were included in the final study (N=6 for CTRL and N=8 for OFCi).

Two subsets of rats, infused only with GRAB-DA2m or GRAB-DA-mut in the VMS and implanted with an optic fiber cannula in that same site, were subjected to the same aforementioned preconditioning protocol for two sessions, after which they underwent a third preconditioning session where the in first three trials of each pairing the cues were presented in the normal order (A→B and C→D), whilst in the last three trials of each pairing the cue order was reversed (B→A and D→C).

### Histological procedures

After completion of the experiment, rats were perfused with chilled phosphate buffer saline (PBS) followed by 4% paraformaldehyde in PBS. The brains were then immersed in 18% sucrose in PBS for at least 24 hours and frozen. The brains were sliced at 40 μm and stained with DAPI (Vectashield-DAPI, Vector Lab, Burlingame, CA), and processed for immunohistochemical detection of green fluorescent protein (Figures 1A and S4B and C). For immunohistochemistry, the brain slices were first blocked in 10% goat serum made in 0.1% Triton X-100/1× PBS and then incubated in anti-GFP antibodies (1/1000, RT, overnight, Mouse anti-GFP, 632381, Takara Bio USA, WI) followed by Alexa-488 secondary antibodies (1/200, RT, 2h, Donkey anti-mouse Alexa- 488, 715-546-150, Jackson, Immunoresearch, PA). Fluorescent microscopy images of the slides were acquired with a BZ-X800 Keyence microscope (Figures 1A and B, and S4B and C).

### Statistical analyses

Statistical analyses were performed in GraphPad Prism (GraphPad Software, San Diego, CA). Error bars in figures denote the standard error of the mean. Effects of experimental variables on behavioral and dopamine signal measures were examined with t-tests if the data was normally distributed or Wilcoxon tests otherwise, as well as repeated-measures 2-way and 3-way ANOVAs combined with Sidak’s or Tukey’s post-hoc tests, respectively. All tests were two-sided. Statistical significance threshold for all tests was set at *P*<0.05.

## Supplementary Information

**Figure S1.**
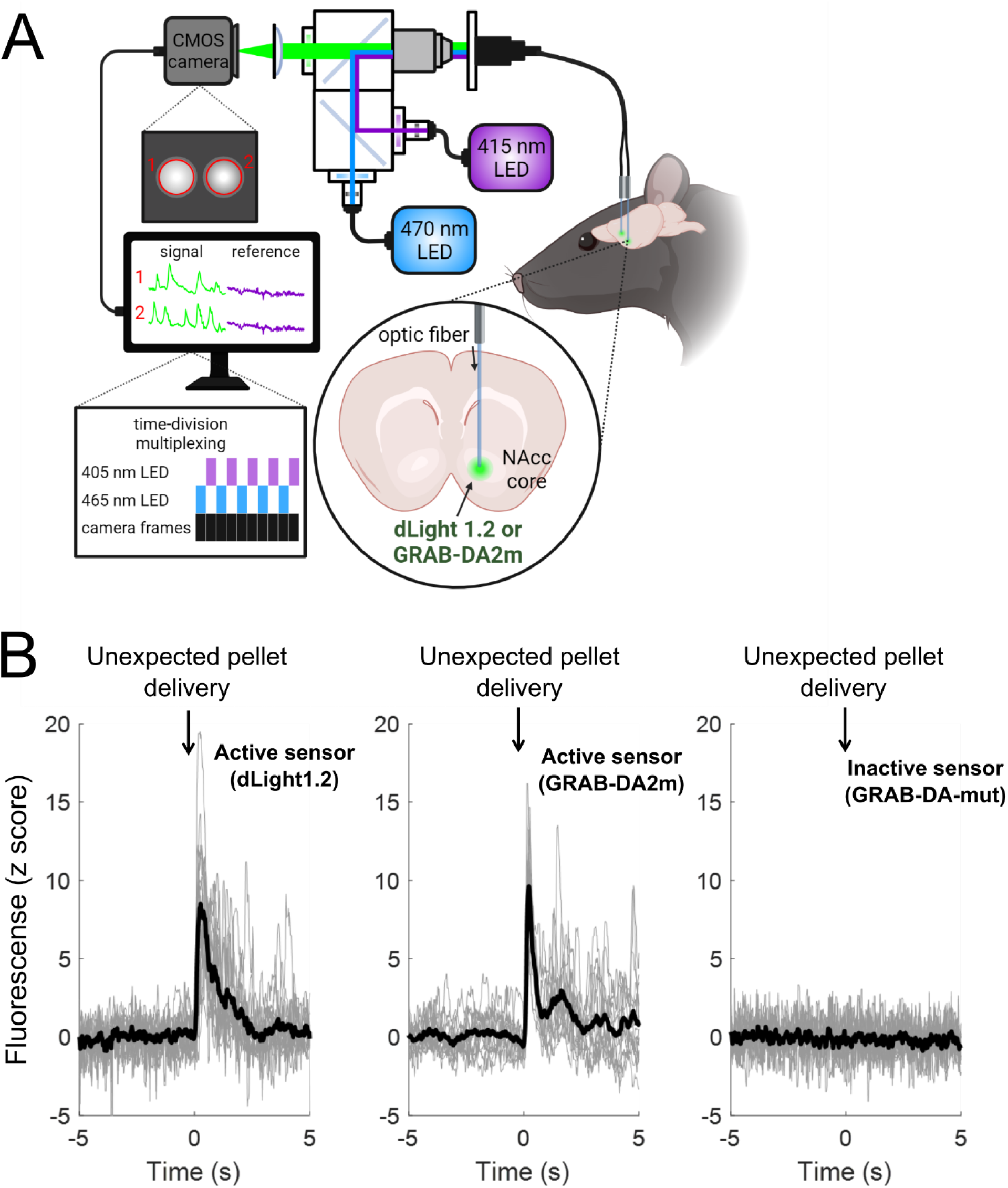
Acquisition and validation of dopamine photometry recordings. **A:** Cartoon schematic of the photometry approach for multi-site recording of dopamine signals in freely moving rats. Rats were implanted with multiple optic fiber cannulas and transfected with AAVs that led to the expression of dopamine sensors in target striatal regions, signals were collected using a Neurophotometrics FP3002 system, and reference (405 nm; ligand-insensitive fluorescence) and active (465) channels are separated using time-division multiplexing. Figure inspired by Martianova et al ^35^ and created with Biorender.com. **B:** Validation of active dopamine sensors (dLight1.2 and GRAB-DA2m) by comparison with a mutated sensor that does not exhibit dopamine-dependent changes in fluorescence ^38^. The panels show data from representative rats, with bold black trace representing mean responses overlayed on gray traces representing 16 individual trials centered around unexpected pellet deliveries in the operant box. Note the clear dopamine responses in the rats transfected with the active sensors, but a complete absence of response in the rats transfected with the mutated control fluorophore.

**Figure S2.**
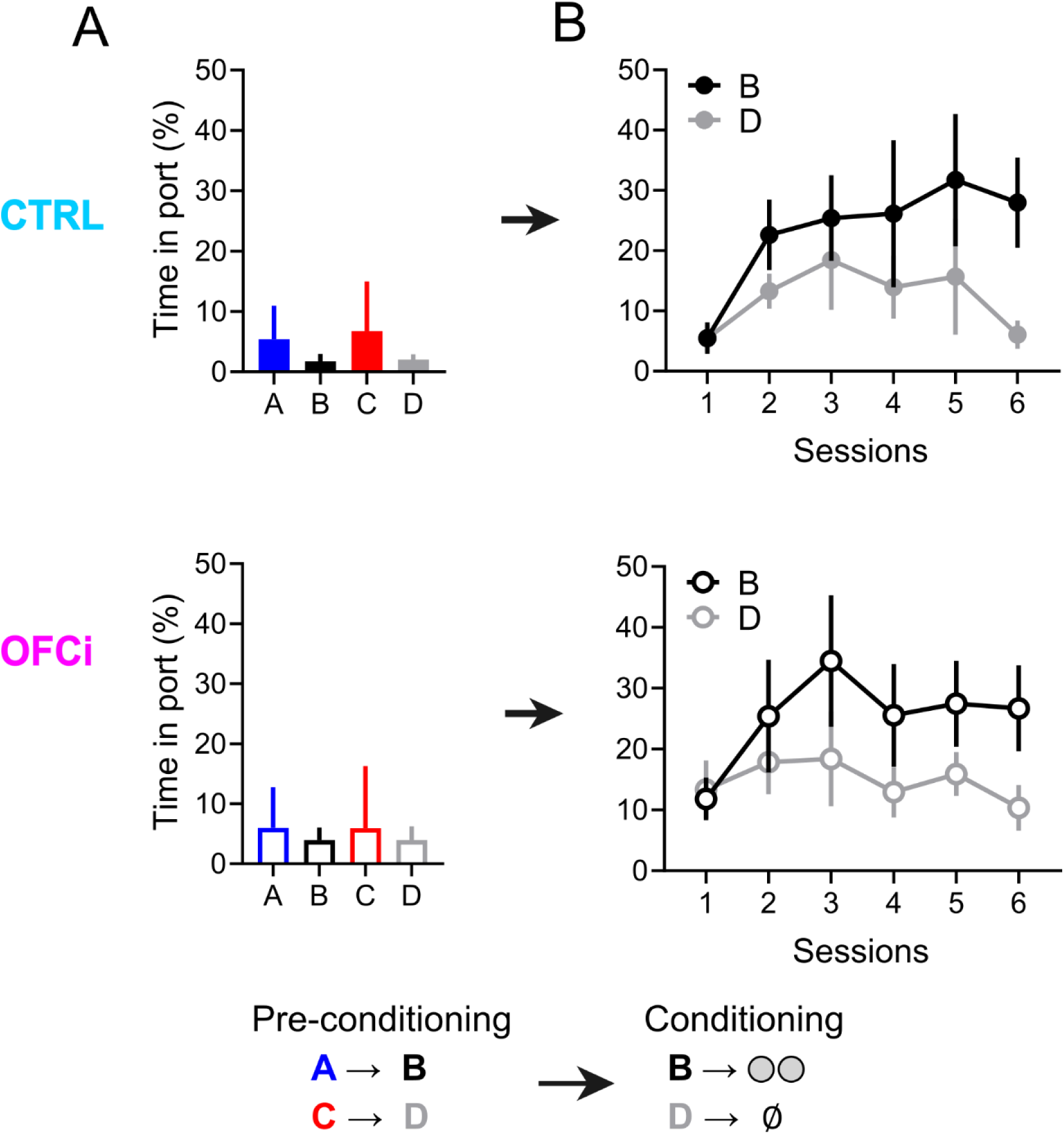
Behavioral responses during preconditioning and conditioning divided by groups. **A:** Percentage of time in port during preconditioning for both CTRL and OFCi rats. **B:** Same metric, but in conditioning. As mentioned in main text, there was no difference in behavior performance between groups in these phases.

**Figure S3.**
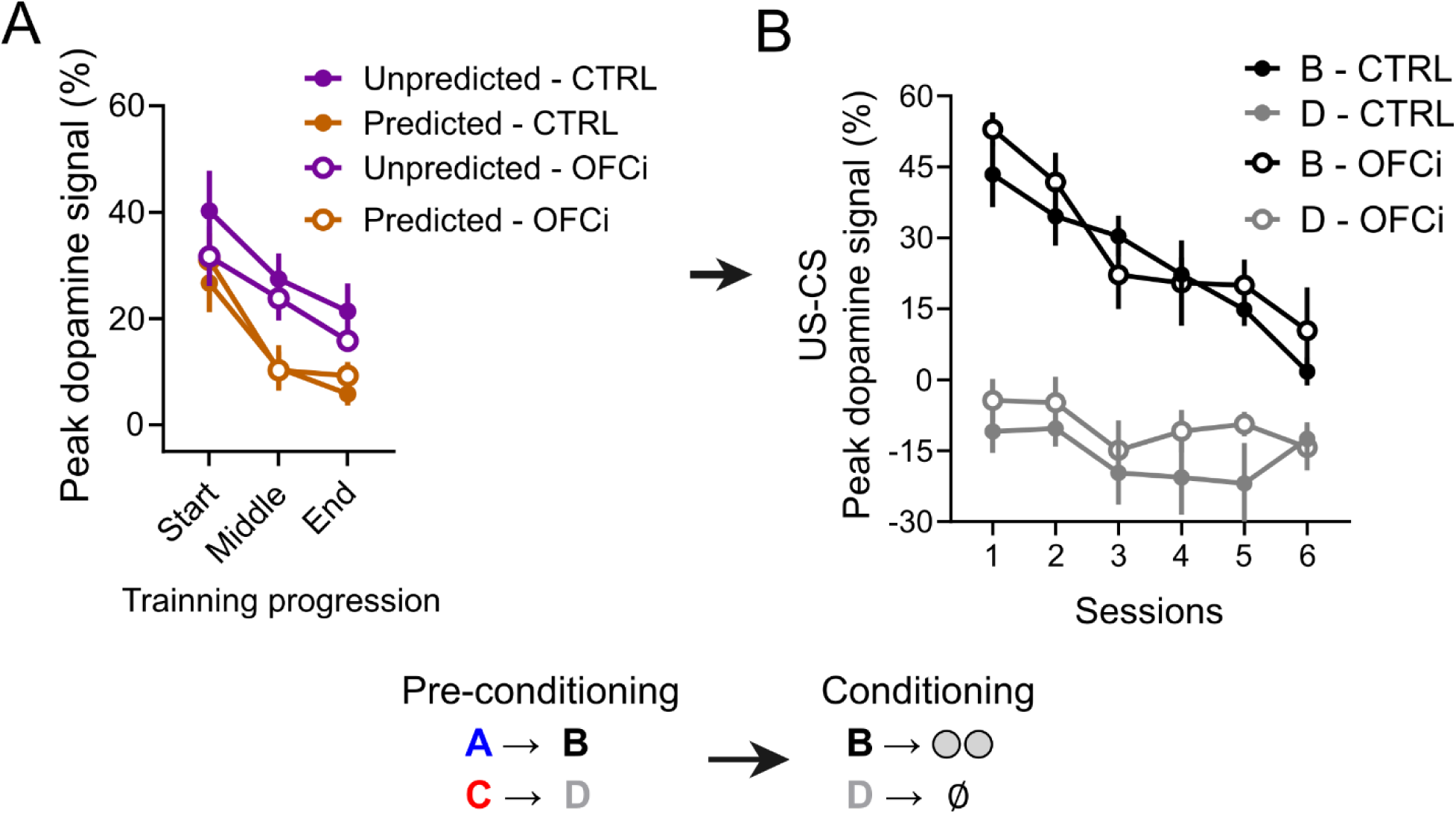
Preconditioning and conditioning dopamine recordings divided by groups. **A:** Peak dopamine responses to predicted and unpredicted cues during preconditioning for both CTRL and OFCi rats. **B:** Difference in peak dopamine responses to the US and CS – a measure of RPE representation – during conditioning sessions. As mentioned in main text, there was no difference in quantified dopamine responses between groups in these phases.

**Figure S4.**
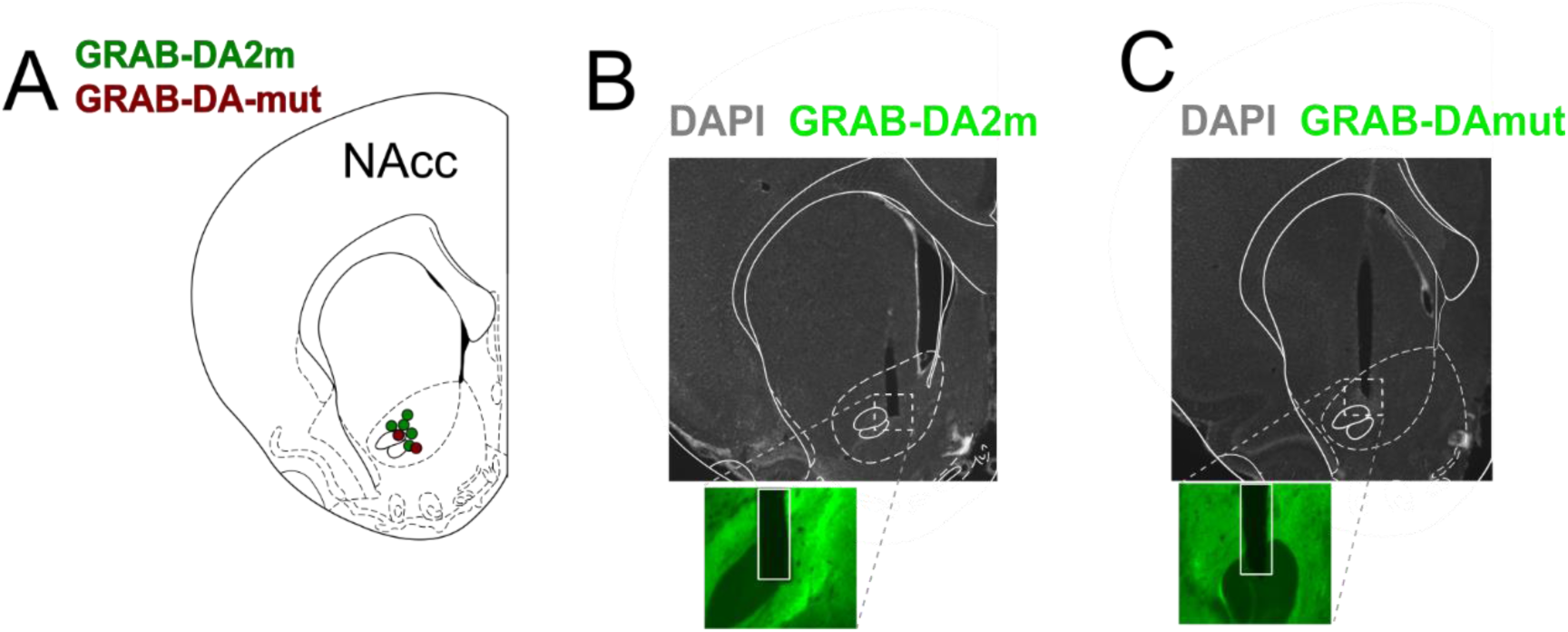
Histological validation of rats transfected with GRAB-DA2m and GRAB-DA-mut. **A:** Schematic representation of fiber tip locations in the brains of the four rats transfected with GRAB-DA2m and used for the preconditioning switch experiment, and for the two rats transfected with GRAB-DA-mut, used as a control for our photometry recordings. **B:** Representative histology of a rat transfected with GRAB- DA2m. C: Same as B, but for a rat transfected with GRAB-DA-mut.

## References

1. Schultz, W., Dayan, P. & Montague, P. R. A Neural Substrate of Prediction and Reward. Science (80-.). 275, 1593–1599 (1997).

2. Tolman, E. C. Cognitive maps in rats and men. Psychol. Rev. 55, 189–208 (1948).

3. Brogden, W. J. Sensory pre-conditioning. J. Exp. Psychol. 25, 323–332 (1939).

4. Costa, K. M. & Schoenbaum, G. Dopamine. Curr. Biol. 32, R817–R824 (2022).

5. Steinberg, E. E. et al. A causal link between prediction errors, dopamine neurons and learning. Nat Neurosci 16, 966–973 (2013).

6. Chang, C. Y. et al. Brief optogenetic inhibition of dopamine neurons mimics endogenous negative reward prediction errors. Nat. Neurosci. 19, 1–8 (2015).

7. Hamilos, A. E. et al. Slowly evolving dopaminergic activity modulates the moment-to- moment probability of reward-related self-timed movements. Elife 10, (2021).

8. Takahashi, Y. K. et al. Dopamine Neurons Respond to Errors in the Prediction of Sensory Features of Expected Rewards. Neuron 95, 1395–1405.e3 (2017).

9. Soares, S., Atallah, B. V. & Paton, J. J. Midbrain dopamine neurons control judgment of time. Science (80-.). 354, 1273–1277 (2016).

10. Gonzalez, L. S. et al. Ventral striatum dopamine release encodes unique properties of visual stimuli in mice. Elife 12, (2023).

11. Sadacca, B. F., Jones, J. L. & Schoenbaum, G. Midbrain dopamine neurons compute inferred and cached value prediction errors in a common framework. Elife 5, e13665 (2016).

12. Papageorgiou, G. K., Baudonnat, M., Cucca, F. & Walton, M. E. Mesolimbic Dopamine Encodes Prediction Errors in a State-Dependent Manner. Cell Rep. 15, 221–228 (2016).

13. Roeper, J. Dissecting the diversity of midbrain dopamine neurons. Trends Neurosci. 36, 336–342 (2013).

14. Farassat, N. et al. In vivo functional diversity of midbrain dopamine neurons within identified axonal projections. Elife 8, (2019).

15. Menegas, W., Babayan, B. M., Uchida, N. & Watabe-Uchida, M. Opposite initialization to novel cues in dopamine signaling in ventral and posterior striatum in mice. Elife 6, 751– 767 (2017).

16. Parker, N. F. et al. Reward and choice encoding in terminals of midbrain dopamine neurons depends on striatal target. Nat. Neurosci. 19, 845–854 (2016).

17. Day, J. J., Roitman, M. F., Wightman, R. M. & Carelli, R. M. Associative learning mediates dynamic shifts in dopamine signaling in the nucleus accumbens. Nat. Neurosci. 10, 1020–1028 (2007).

18. Kutlu, M. G. et al. Dopamine release in the nucleus accumbens core signals perceived saliency. Curr. Biol. 31, 4748–4761.e8 (2021).

19. Kutlu, M. G. et al. Dopamine signaling in the nucleus accumbens core mediates latent inhibition. Nat. Neurosci. 25, 1071–1081 (2022).

20. Stalnaker, T. A. et al. Dopamine neuron ensembles signal the content of sensory prediction errors. Elife 8, (2019).

21. Keiflin, R., Pribut, H. J., Shah, N. B. & Janak, P. H. Ventral Tegmental Dopamine Neurons Participate in Reward Identity Predictions. Curr. Biol. 29, 93–103.e3 (2019).

22. Chang, C. Y., Gardner, M., Di Tillio, M. G. & Schoenbaum, G. Optogenetic Blockade of Dopamine Transients Prevents Learning Induced by Changes in Reward Features. Curr. Biol. 27, 3480–3486.e3 (2017).

23. Sharpe, M. J. et al. Dopamine transients are sufficient and necessary for acquisition of model-based associations. Nat. Neurosci. 20, 735–742 (2017).

24. Takahashi, Y. K. et al. Dopaminergic prediction errors in the ventral tegmental area reflect a multithreaded predictive model. Nat. Neurosci. 2023 265 26, 830–839 (2023).

25. Millidge, B., Song, Y., Lak, A., Walton, M. E. & Bogacz, R. Reward-Bases: Dopaminergic Mechanisms for Adaptive Acquisition of Multiple Reward Types. bioRxiv 2023.05.09.540067 (2023) doi:10.1101/2023.05.09.540067.

26. 26. Lee, R. S., Engelhard, B., Witten, I. B. & Daw, N. D. A vector reward prediction error model explains dopaminergic heterogeneity. bioRxiv 2022.02.28.482379 (2022) doi:10.1101/2022.02.28.482379.

27. Wang, Y., Toyoshima, O., Kunimatsu, J., Yamada, H. & Matsumoto, M. Tonic firing mode of midbrain dopamine neurons continuously tracks reward values changing moment-by- moment. Elife 10, (2021).

28. Schultz, W. Recent advances in understanding the role of phasic dopamine activity. F1000Research 2019 81680 8, 1680 (2019).

29. Gardner, M. P. H., Schoenbaum, G. & Gershman, S. J. Rethinking dopamine as generalized prediction error. Proc. R. Soc. B Biol. Sci. 285, 20181645 (2018).

30. Langdon, A. J., Sharpe, M. J., Schoenbaum, G. & Niv, Y. Model-based predictions for dopamine. Curr. Opin. Neurobiol. 49, 1–7 (2018).

31. Berridge, K. C. The debate over dopamine’s role in reward: the case for incentive salience. Psychopharmacology (Berl). 191, 391–431 (2007).

32. Lau, B., Monteiro, T. & Paton, J. J. The many worlds hypothesis of dopamine prediction error: implications of a parallel circuit architecture in the basal ganglia. Curr. Opin. Neurobiol. 46, 241–247 (2017).

33. Corlett, P. R., Mollick, J. A. & Kober, H. Meta-analysis of human prediction error for incentives, perception, cognition, and action. Neuropsychopharmacology 47, 1339–1349 (2022).

34. Jones, J. L. et al. Orbitofrontal cortex supports behavior and learning using inferred but not cached values. Science (80-.). 338, 953–956 (2012).

35. Martianova, E., Aronson, S. & Proulx, C. D. Multi-Fiber Photometry to Record Neural Activity in Freely-Moving Animals. JoVE (Journal Vis. Exp. 2019, e60278 (2019).

36. Patriarchi, T. et al. Ultrafast neuronal imaging of dopamine dynamics with designed genetically encoded sensors. Science (80-.). 360, (2018).

37. Lopes, G. et al. Bonsai: an event-based framework for processing and controlling data streams. Front. Neuroinform. 9, (2015).

38. Sun, F. et al. Next-generation GRAB sensors for monitoring dopaminergic activity in vivo. Nat. Methods 2020 1711 17, 1156–1166 (2020).

39. Costa, K. M., Sengupta, A. & Schoenbaum, G. The orbitofrontal cortex is necessary for learning to ignore. Curr. Biol. 31, 2652–2657.e3 (2021).

40. Costa, K. M. et al. The role of the lateral orbitofrontal cortex in creating cognitive maps. Nat. Neurosci. 2022 261 26, 107–115 (2022).

41. Bonaventura, J. et al. High-potency ligands for DREADD imaging and activation in rodents and monkeys. Nat. Commun. 10, 1–12 (2019).

42. Gardner, M. P. H., Conroy, J. S., Shaham, M. H., Styer, C. V. & Schoenbaum, G. Lateral Orbitofrontal Inactivation Dissociates Devaluation-Sensitive Behavior and Economic Choice. Neuron 96, 1192–1203.e4 (2017).

43. Kakade, S. & Dayan, P. Dopamine: Generalization and bonuses. Neural Networks 15, 549–559 (2002).

44. Schultz, W. Dopamine reward prediction-error signalling: a two-component response. Nat. Rev. Neurosci. 17, 183–195 (2016).

45. Bunzeck, N. & Düzel, E. Absolute Coding of Stimulus Novelty in the Human Substantia Nigra/VTA. Neuron 51, 369–379 (2006).

46. Morrens, J., Aydin, Ç., Janse van Rensburg, A., Esquivelzeta Rabell, J. & Haesler, S. Cue-Evoked Dopamine Promotes Conditioned Responding during Learning. Neuron 106, 142–153.e7 (2020).

47. Haber, S. N., Fudge, J. L. & McFarland, N. R. Striatonigrostriatal pathways in primates form an ascending spiral from the shell to the dorsolateral striatum. J. Neurosci. 20, 2369–82 (2000).

48. Willuhn, I., Burgeno, L. M., Everitt, B. J. & Phillips, P. E. M. Hierarchical recruitment of phasic dopamine signaling in the striatum during the progression of cocaine use. Proc. Natl. Acad. Sci. U. S. A. 109, 20703–8 (2012).

49. Mohebi, A. et al. Dissociable dopamine dynamics for learning and motivation. Nat. 2019 5707759 570, 65–70 (2019).

50. Hart, E. E., Sharpe, M. J., Gardner, M. P. & Schoenbaum, G. Responding to preconditioned cues is devaluation sensitive and requires orbitofrontal cortex during cue- cue learning. Elife 9, (2020).

51. Engelhard, B. et al. Specialized coding of sensory, motor and cognitive variables in VTA dopamine neurons. Nat. 2019 5707762 570, 509–513 (2019).

52. Kidd, C. & Hayden, B. Y. The Psychology and Neuroscience of Curiosity. Neuron 88, 449–460 (2015).

53. Gottlieb, J., Oudeyer, P. Y., Lopes, M. & Baranes, A. Information-seeking, curiosity, and attention: computational and neural mechanisms. Trends Cogn. Sci. 17, 585–593 (2013).

54. Jeong, H. et al. Mesolimbic dopamine release conveys causal associations. Science (80-.). 378, (2022).

55. Kobayashi, S. & Schultz, W. Reward contexts extend dopamine signals to unrewarded stimuli. Curr. Biol. 24, 56–62 (2014).

56. Sharpe, M. J., Batchelor, H. M. & Schoenbaum, G. Preconditioned cues have no value. Elife 6, (2017).

57. Friston, K. The free-energy principle: a unified brain theory? Nat. Rev. Neurosci. 2010 112 11, 127–138 (2010).

58. Wilson, R. C., Takahashi, Y. K., Schoenbaum, G. & Niv, Y. Orbitofrontal cortex as a cognitive map of task space. Neuron 81, 267–279 (2014).

59. Liu, C. et al. An action potential initiation mechanism in distal axons for the control of dopamine release. Science (80-.). 375, 1378–1385 (2022).

60. Kramer, P. F. et al. Synaptic-like axo-axonal transmission from striatal cholinergic interneurons onto dopaminergic fibers. Neuron 110, 2949–2960.e4 (2022).

61. Zhang, H. & Sulzer, D. Glutamate Spillover in the Striatum Depresses Dopaminergic Transmission by Activating Group I Metabotropic Glutamate Receptors. J. Neurosci. 23, 10585–10592 (2003).

